# IFN-independent G0 arrest and SAMHD1 activation following TLR4 activation in macrophages

**DOI:** 10.1101/741124

**Authors:** P. Mlcochova, H. Winstone, L. Zuliani-Alvarez, R.K. Gupta

## Abstract

Monocyte-derived macrophages mostly reside in a resting, G0 state, expressing high levels of dephosphorylated, active SAMHD1. We have previously shown that macrophages can re-enter the cell cycle without division, into a G1-like state. This cell cycle re-entry is accompanied by phosphorylation of the dNTP hydrolase/ antiviral restriction factor SAMHD1 at T592 by the cyclin-dependent kinase CDK1. HIV-1 successfully infects macrophages in G1 through exploiting this naturally occurring window of opportunity where SAMHD1 antiviral activity is de-activated.

Here we demonstrate for the first time that LPS activation of the pathogen associated molecular pattern (PAMP) receptor TLR4 induces G0 arrest in human macrophages. We show this G0 arrest is MyD88-independent and therefore NFkB independent. Furthermore, the effect of TLR4 activation on cell cycle is regulated by (a) the canonical IFN-dependent pathway following TBK1 activation and IRF3 translocation and (b) an IFN-independent pathway that occurs prior to TBK1 activation, and that is accompanied by CDK1 downregulation, p21 upregulation and SAMHD1 dephosphorylation at T592. Furthermore, we show by siRNA knockdown of SAMHD1 that the interferon independent pathway activated by TLR4 is able to potently block HIV-1 infection in macrophages specifically via SAMHD1. Finally, ingestion of whole E. Coli and TLR4 activation by macrophages also activates SAMHD1 via the interferon independent pathway.

Together, these data demonstrate that macrophages can rapidly activate an intrinsic cell arrest and anti-viral state by activation of TLR4 prior to IFN secretion, thereby highlighting the importance of cell cycle regulation as a response to danger signals in human macrophages. Interferon independent activation of SAMHD1 by TLR4 represents a novel mechanism for limiting the HIV-1 reservoir size and should be considered for host-directed therapeutic approaches that may contribute to curative interventions.

## Introduction

Macrophages are the first line of defence against invading pathogens, sensing through pathogen recognition receptors (PRRs) and initiating innate and adaptive responses. The most studied PRRs are Toll-like receptors (TLRs), expressed in monocytes, macrophages and dendritic cells. They play a fundamental role in recognition of pathogen-associated molecular patterns expressed on infectious agents, and subsequently initiate a series of inflammatory events that depend upon the MyD88 and/or TRIF signalling pathways ^1^. Amongst TLRs, TLR4 acts as a receptor for lipopolysaccharide (LPS), a component from the wall of gram-negative bacteria ^2^. Once activated it triggers a well characterised downstream-signalling cascade involving multiple signalling components, culminating in the activation of transcription factors, such as NFkB and IRFs, which, in turn induce various immune and pro-inflammatory genes ^3^.

Many studies in the past have shown that LPS has a potent inhibitory activity against HIV-1 infection ^4–10^. LPS has been shown to down-regulate receptors for HIV-1 entry and impair early steps of viral life cycle ^5, 11, 12^. The mediators of HIV-1 suppression by LPS-stimulated MDM are mostly secreted 1)-chemokines and interferons (IFN) ^4, 7, 9, 11^. However, some data suggest that IFN release by LPS-stimulated macrophages/dendritic cells might not be the main mediator of HIV-1 suppression ^9, 10^. The diverse data on effect of LPS in HIV-1 infection suggests multiple anti-HIV mechanisms, consistent with up-regulation of an array of HIV-1 restriction factors ^7, 10, 13^ by IFN in LPS-stimulated macrophages and dendritic cells.

Terminally differentiated myeloid cells and resting T cells express SAMHD1, a deoxynucleotide-triphosphate (dNTP) hydrolase which restricts HIV-1 reverse transcription (RT) through decreasing levels of dNTPs ^14, 15^. SAMHD1 phosphorylation at position T592 mediated by cyclin-dependent kinases CDK1/2 ^16, 17^ impairs its dNTP hydrolase activity in actively dividing cells and allows viral DNA synthesis to occur ^16, 18^. SAMHD1 in its dephosphorylated form is active against HIV-1 and known to block infection ^16, 17, 19, 20^.

Our group recently showed that SAMHD1 restriction capacity can be manipulated in monocyte-derived macrophages (MDM). We demonstrated that macrophages, cells normally residing in a G0/terminally differentiated state can re-enter the cell cycle into a G1-like phase, expressing certain cellular cell cycle factors, including CDK1 that is known to phosphorylate and deactivate the antiviral activity of SAMHD1 ^19, 21^.

Type-I interferons are known to lead to dephosphorylation/activation of SAMHD1 in MDM ^16, 22^. Whilst it has been proposed that LPS-stimulated macrophages can mediate HIV-1 restriction through IFN secretion, the role of SAMHD1 has not been established.

Here we show that TLR4 activation by bacterial LPS can mediate HIV-1 inhibition through regulation of SAMHD1 in an IFN-independent manner, as well as via the canonical IFN dependent pathway. We show that this IFN-independent pathway is MyD88-independent, occurring prior to TBK1 activation, and resulting in p21 upregulation and G0 arrest with SAMHD1 dephosphorylation. This is the first demonstration that TLR4 activation can directly induce G0 arrest in human macrophages, adding to growing evidence that G0 arrest is a conserved and important response to danger signals even in cells classically viewed as being terminally differentiated.

## Results

### Interferon-independent SAMHD1 dephosphorylation and HIV-1 blockade in macrophages following TLR4 activation

We used human monocyte derived macrophages (MDM) to study the effect of LPS on permissivity to HIV-1 infection. Macrophages were exposed to 10ng/ml LPS for 18h and infected with VSV-G pseudotyped HIV-1 expressing GFP. LPS treatment completely inhibited HIV-1 infection of MDM, accompanied by activation/dephosphorylation of SAMHD1 at T592 (Fig.1A,B).

**FIGURE 1:**
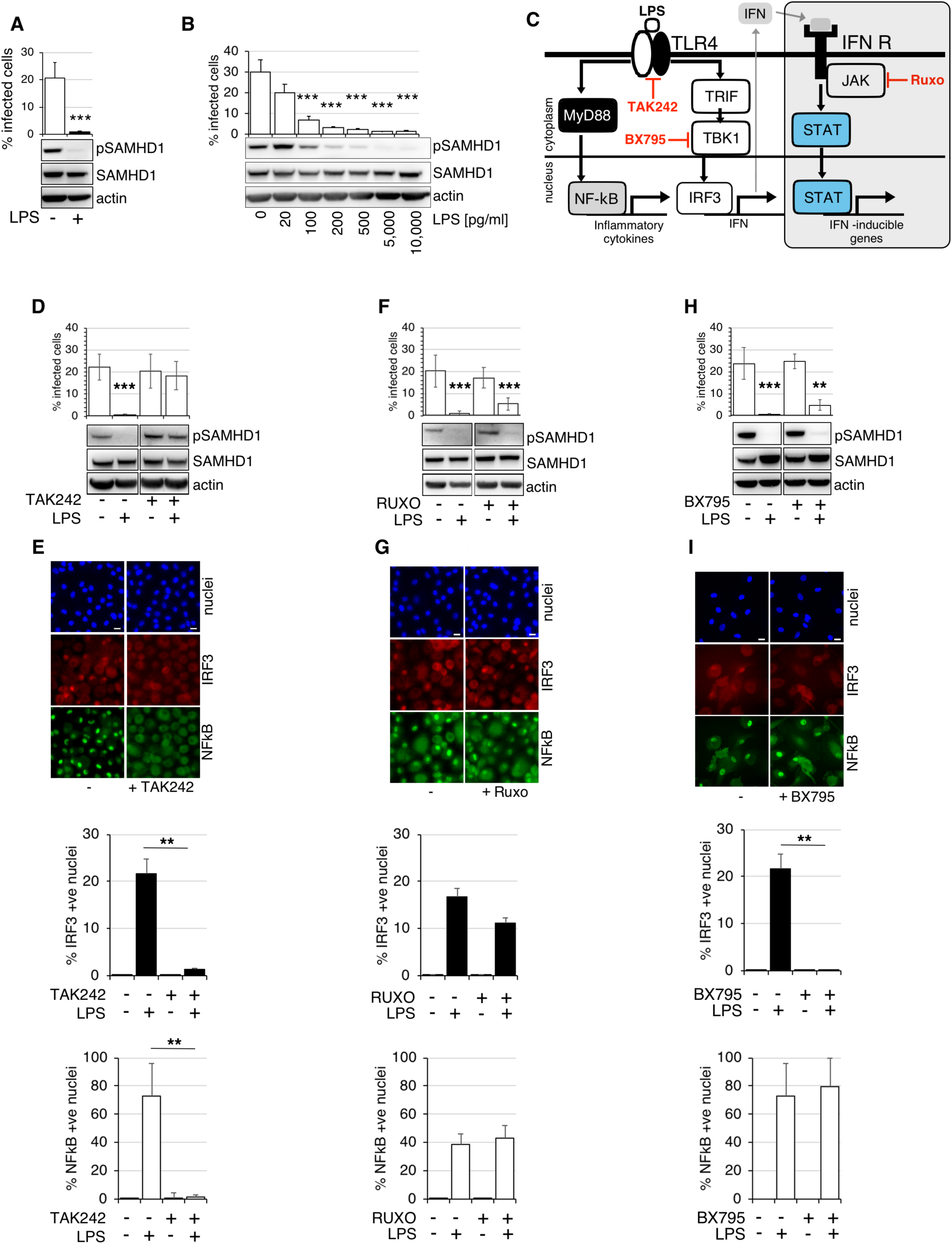
TLR4 activation dephosphorylates SAMHD1 and blocks HIV-1 infection. A. MDM were treated by LPS for 18h and infected by VSV-G pseudotyped HIV-% infected cells were determined 48h post-infection. (*n* = 6, mean ± s.e.m.; ***P-value≤*0.001*, paired *t*-test). Cells from a representative donor were used for immunoblotting. B. MDM were treated with increasing concentration of LPS 18h before infection. Cells were infected by VSV-G pseudotyped HIV-1 and % infected cells were determined 48h post-infection. (*n* = 3, mean ± s.e.m.; ***P-value≤*0.001,* paired *t*-test). Cells from a representative donor were used for immunoblotting. C. A simplified diagram of macrophage activation mediated by TLR4 signalling in response to LPS. LPS activates both MyD88-dependent and independent signalling pathways. TAK242, an inhibitor of TLR4 signalling; BX795, an inhibitor of TBK1, suppresses IFN signalling; RUXO, ruxolitinib, inhibitor of JAK1/2 kinase, suppresses IFN signalling. D, F, H. MDM were treated with inhibitors for 6h (D) TLR4 inhibitor TAK242, (F) JAK1/2 inhibitor RUXO, or 2h (H) TBK1 inhibitor BX795 before addition of LPS. Cells were infected by VSV-G pseudotyped HIV-1 18h later. % infected cells were determined 48h post-infection. (*n* = 3, mean ± s.e.m.; ***P-value≤*0.001,**P*-value ≤ 0.01, paired *t*-test). Cells from a representative donor were used for immunoblotting. E,G,I. IRF3/NFkB translocation assay. Cells were exposed to LPS in the absence or presence of (E) TAK242, (G) RUXO (I) BX795 and 2h later stained for IRF3/NFkB. % of cells with nuclear staining were determined. (*n* = 3, mean ± s.e.m.; *\*\*P-*value ≤ 0.01, paired *t*-test).

We next confirmed that the LPS effect on HIV-1 infection was directly through TLR4 activation (Fig.1C). We used the TLR4 inhibitor TAK242 that in the presence of LPS was able to rescue HIV-1 infection of MDM and prevent SAMHD1 activation/dephosphorylation (Fig.1D). TAK242 also prevented translocation of NFkB and IRF3 into nucleus in the presence of LPS, confirming TLR4 inhibition (Fig. 1C,E).

TLR4 activation stimulates production of interferons (IFN). In order to understand if IFN plays role in SAMHD1 activation/dephosphorylation after TLR4 activation as recently published ^16, 22^, we used the JAK1/2 inhibitor Ruxolitinib (RUXO) that inhibits IFN signalling (Fig.1C)^23^. The inhibitor of IFN signalling could not restore infection and failed to reverse SAMHD1 phosphorylation at T592, suggesting that regulation of SAMHD1 phosphorylation is largely independent of IFN (Fig.1F). Importantly, RUXO had no effect on nuclear IRF3/NFkB translocation following LPS addition (Fig.1G).

Next, we used the TBK1 inhibitor BX795 (Fig.1C,H,I) that blocks the phosphorylation, nuclear translocation, and transcriptional activity of IRF3 and production of interferon in response to TLR3 and TLR4 agonists ^24^ (Fig.1I). BX795 treatment also failed to restore infection and SAMHD1 phosphorylation after LPS addition (Fig.1H), confirming that SAMHD1 dephosphorylation/activation is IFN-independent and is mediated upstream from TBK1.

### SAMHD1 activation following TLR4 engagement is MyD88 independent

Tenascin-C (TNC) is an extracellular matrix protein which expression is rapidly induced at the site of infection or injury where it triggers inflammation by activating TLR4 in different cells including macrophages ^25^. This TLR4 activation is specifically mediated through the MyD88-dependent pathway ^26^ (Fig. 2A). TNC effectively activated the MyD88 dependent pathway in MDM, which was confirmed by detectable translocation of NFkB but absence of nuclear IRF3 (Fig. 2B) and by production of cytokines into culture media after treatment with LPS and TNC (Fig.2C). Importantly, when MDM were treated with TNC, no SAMHD1 dephosphorylation or HIV-1 inhibition was detected (Fig. 2D). From these experiments we conclude that LPS activates TLR4 and blocks HIV-1 infection in a MyD88-independent pathway, culminating in dephosphorylation of SAMHD1.

**FIGURE 2:**
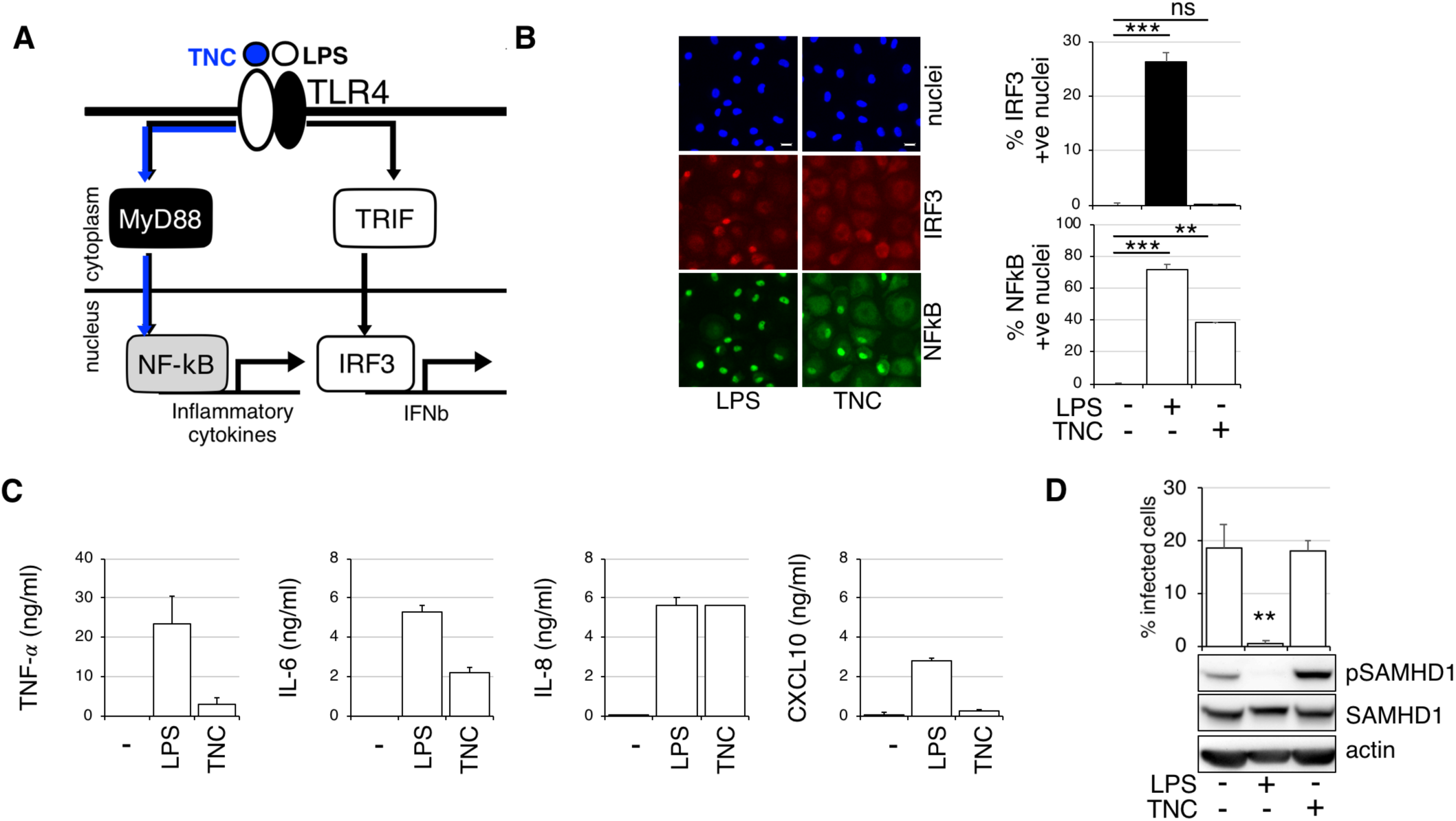
TLR4 activation of SAMHD1 is MyD88-independent. A. Diagram of TLR4 activation. LPS activates both MyD88-dependent and independent signalling pathways. Tenascin-C (TNC) activates only MyD88-dependent pathway leading to NFkB translocation to nucleus. B. IRF3/NFkB translocation assay. Cells were exposed to TNC and LPS and 2h later stained for IRF3/NFkB. % of cells with nuclear staining were determined. (*n* = 3, mean ± s.e.m.; ***P-value≤*0.001,**P*-value ≤ 0.01, (ns) non-significant, paired *t*-test). C. MDM were treated with LPS and TNC and cytokines were measured by ELISA in culture media 24h later. D. MDM were treated with TNC and LPS. Cells were infected by VSV-G pseudotyped HIV-1 18h later. % infected cells were determined 48h post-infection. (*n* = 3, mean ± s.e.m.; *\*\*P*-value ≤ 0.01, paired *t*-test). Cells from a representative donor were used for immunoblotting.

### TLR4 engagement results in interferon-independent G0 arrest in macrophages

We have shown previously that macrophage entry to a G1-like state is accompanied by an increase in certain cell cycle associated proteins such as MCM2 and CDK1, as well as phosphorylation of SAMHD1 at T592 that confers increased susceptibility to HIV-1 infection ^19, 21^. LPS treatment resulted in a decrease of MCM2 expression and as expected, SAMHD1 dephosphorylation at T592 (Fig.3A, S1A,B). These results suggest that LPS treatment led to cell arrest in macrophages where SAMHD1 is activated and can block HIV-1 infection as shown in Fig.3A,B. Crucially, the TLR4 inhibitor TAK242 completely reversed the cell arrest and SAMHD1 phosphorylation changes, restoring HIV-1 infection. As expected, blocking IFN signalling in macrophages by treatment with RUXO or TBK1 inhibitor BX795 could not restore cell cycle changes, SAMHD1 phosphorylation and HIV-1 infection (Fig.3A,B). Remarkably, addition of TNC, molecule that activates TLR4 exclusively via the MyD88-dependent signalling pathway, had no effect on cell cycle in MDM (Fig.3C). These data strongly suggest that HIV-1 restriction is mediated through G0 arrest that is uncoupled from IRF3 signalling.

**FIGURE 3:**
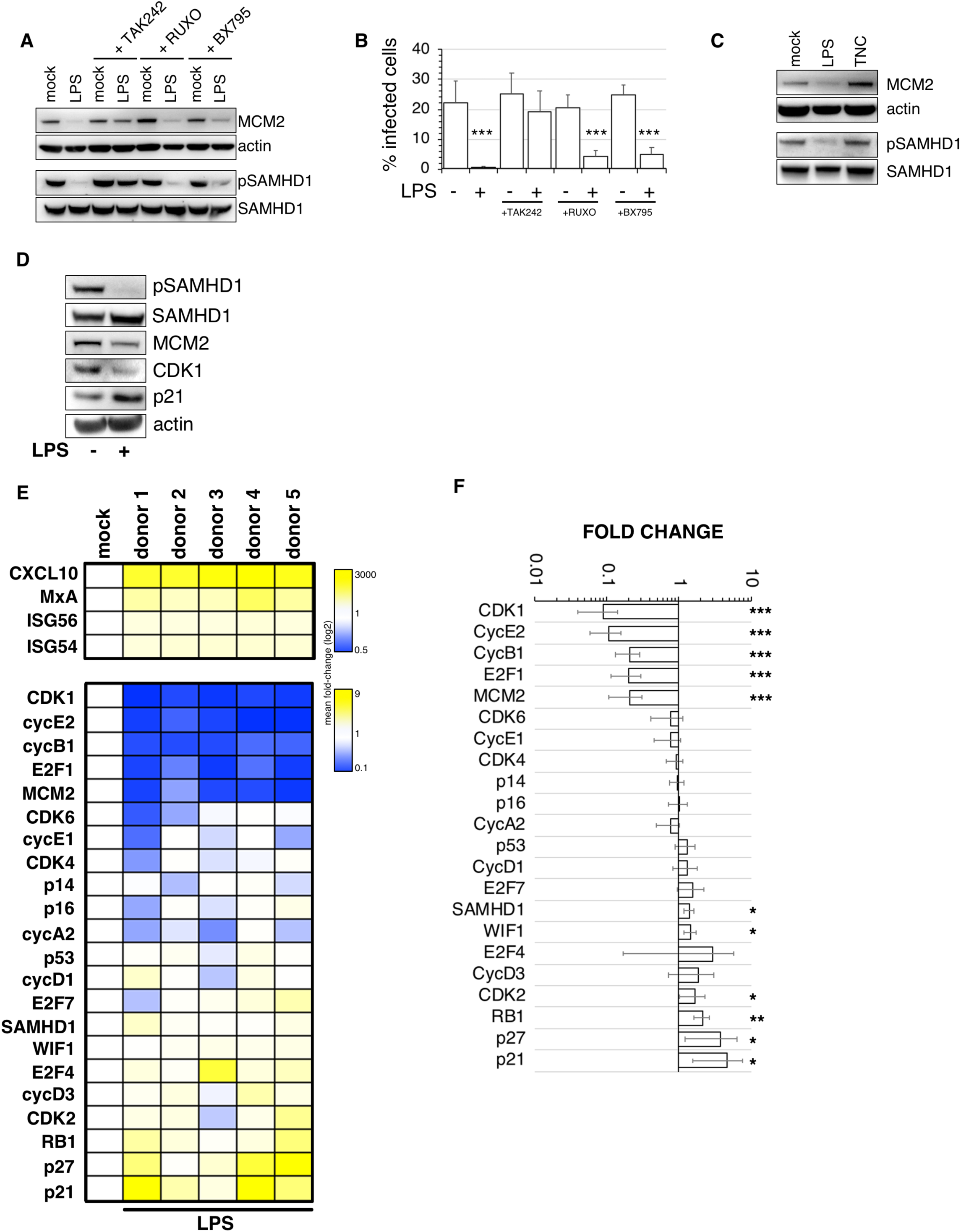
TLR4 activation induces G0 arrest in human MDM. A. MDM were treated with TAK242, RUXO, and BX795 6h before addition of LPS. Cell lysates were prepared 18h later to detect changes to cells cycle and SAMHD1 phosphorylation. MCM2, a marker of cell cycle is expressed during cell cycle phases and completely absent in G0/quiescent state. B. MDM were treated with TAK242, RUXO, and BX795 6h before addition of LPS. Cells were infected by VSV-G pseudotyped HIV-1 18h later. % infected cells were determined 48h post-infection. (*n* = 3, mean ± s.e.m.; ***P-value≤*0.001*, paired *t*-test). C. MDM were treated with TNC and LPS. Cell lysates were prepared 18h later to detect changes to cells cycle and SAMHD1 phosphorylation. MCM2, a marker of cell cycle is expressed during cell cycle phases and completely absent in G0/quiescent state. D. MDM were treated by LPS and cell lysates were prepared 18h later to detect changes in cell cycle associated proteins. E. A heat map presents differential gene expression patterns of cell cycle associated transcripts in MDM from 5 different donors treated with LPS for 18h. The colour scale bar corresponds to log-fold expression. F. Relative expression levels (fold change) of cell cycle associated transcripts. (*n* = 6, mean ± s.e.m.; ***P-value≤*0.001, **P*-value ≤ 0.01, *\*P*-value ≤ 0.1, paired *t*-test).

We hypothesised that the LPS induced G0 arrest would be regulated by expression of negative cell cycle regulators such as p21 or p27. Indeed, immunoblotting confirmed that the decrease in CDK1 and MCM2 after LPS treatment was accompanied by increased p21 levels (Fig.3D). As we were unable to detect p27 expression in immunoblot we next sought to further characterise the cell cycle program changes triggered by LPS in MDM, using a panel of cell cycle associated transcripts (Fig.3E,F). Statistically significant decreases were observed in transcripts associated with cell cycle progression: CDK1, cyclin E2, cyclin B1, E2F1, MCM2 transcripts compared to the untreated control (set to 1). A small but significant increase was observed for SAMHD1 and WIF1 transcripts, and also for transcripts associated with G0 arrest or G0 state: CDK2, RB1, p27 and p21. These data show that the macrophage cell cycle program is impacted by TLR4 engagement, and are the first evidence of G0 arrest induced by LPS treatment in primary human macrophages.

### Type-I Interferon impacts cell cycle in human macrophages

TLR4 activation of a TRIF dependent pathway leads to IRF3 translocation into the nucleus and expression of interferon (IFN). IFN as a cause of G0 arrest has been reported in myeloid cells from mice and murine cell lines as well as in human cell lines ^27–29^, monocytes and T-cells ^30^. The effect of IFN on cell cycle states in primary human macrophages has not been reported.

We therefore examined the effect of blocking IFN signalling after TLR4 activation as well as after addition of exogenous IFNβ) (Fig.4). Ruxolitinib (RUXO), an inhibitor of JAK kinases and IFN signalling, was used to treat MDM 6h before addition of LPS or IFNβ). As expected RUXO completely blocked expression of selected ISG: CXCL10, MxA, ISG 54 and 56 after both LPS and IFNβ) treatment, confirming that IFN signalling is inhibited in both conditions (Fig.4A).

**FIGURE 4:**
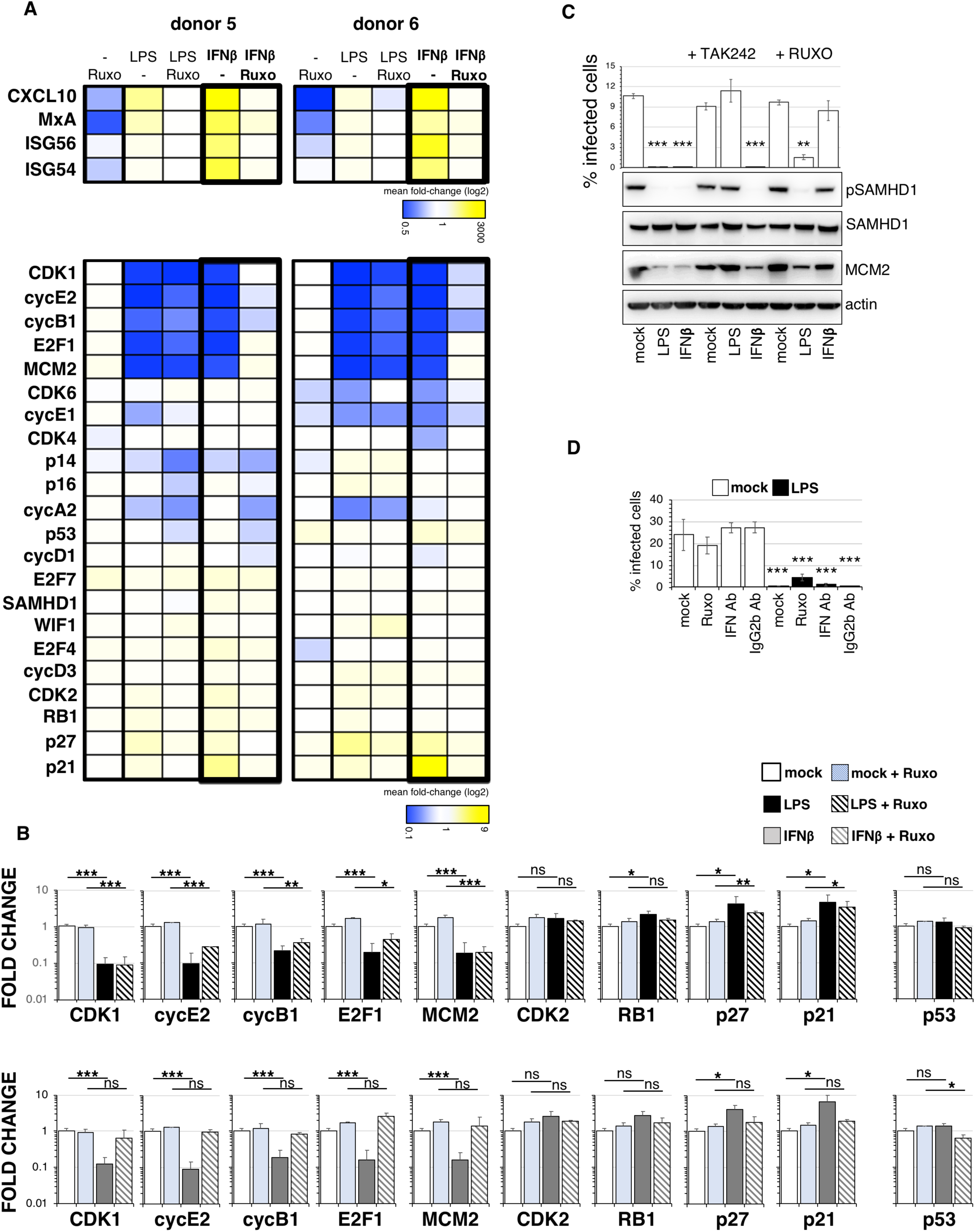
LPS mediated cell cycle regulation in human MDM is both interferon-dependent and independent. A. A heat map presents differential gene expression patterns of cell cycle associated transcripts in MDM treated with LPS or IFNβ) in the presence or absence of RUXO. The colour scale bar corresponds to log-fold expression. B. Relative expression levels (fold changes) of statistically significantly changed cell cycle associated transcripts after LPS or IFNβ) treatment in the presence or absence of RUXO. (*n* = 4, mean ± s.e.m.; ***P-value≤*0.001, **P*-value ≤ 0.01, *\*P*-value ≤ 0.1, (ns) non-significant, paired *t*-test). C. MDM were treated with TAK242 and RUXO 6h before addition of LPS and interferon b (IFNβ)). Cells were infected by VSV-G pseudotyped HIV-1 18h later. % infected cells were determined 48h post-infection. (*n* = 3, mean ± s.e.m.; ***P-value≤*0.001, **P*-value ≤ 0.01, paired *t*-test). Cells from a representative donor were used for immunoblotting. D. MDM were exposed to anti-IFN Ab/IgG2b non-specific Ab and treated with LPS. Cells were infected by VSV-G pseudotyped HIV-1 18h later. % infected cells were determined 48h post-infection. (*n* = 3, mean ± s.e.m.; *\*\*\*P*-value ≤ 0.01, paired *t*-test).

While G0 arrest was observed after both LPS and exogenous IFNβ) addition, based on expression levels of cell cycle associated transcripts, RUXO was unable to rescue G0 arrest caused by LPS/TLR4 activation but completely rescued G0 arrest after addition of exogenous IFNβ) (Fig.4A,B). This confirms redundancy of pathways activated by TLR4 that lead to G0 arrest. In concordance with these results RUXO failed to restore HIV-1 infection from the effect of LPS but completely rescued HIV-1 infection after exposure to exogenous IFNβ) (Fig.4C). These data were confirmed by using an IFN receptor antibody instead of RUXO (Fig.4D). As a control the TLR4 inhibitor TAK242 rescued cell cycle and HIV-1 infection after LPS treatment. As expected, TAK242 could not rescue either after IFNβ) treatment.

We conclude existence of two independent pathways that are responsible for G0 arrest in MDM, both of which are able to potently block HIV-1 infection. G0 arrest being the first early block to HIV-1 infection while IFN production representing a second wave.

### SAMHD1 is directly responsible for the interferon independent HIV-1 blockade following TLR4 activation

We have shown previously that the restriction of HIV-1 infection in G0 MDM can be completely lifted by SAMHD1 depletion ^19^. Our experimental system here involves use of MDM predominantly in G1, where SAMHD1 is deactivated/phosphorylated. To confirm that SAMHD1 dephosphorylation/activation is responsible for block to HIV-1 infection after TLR4 activation we employed SAMHD1 knock-down (KD) (Fig.5).

**FIGURE 5:**
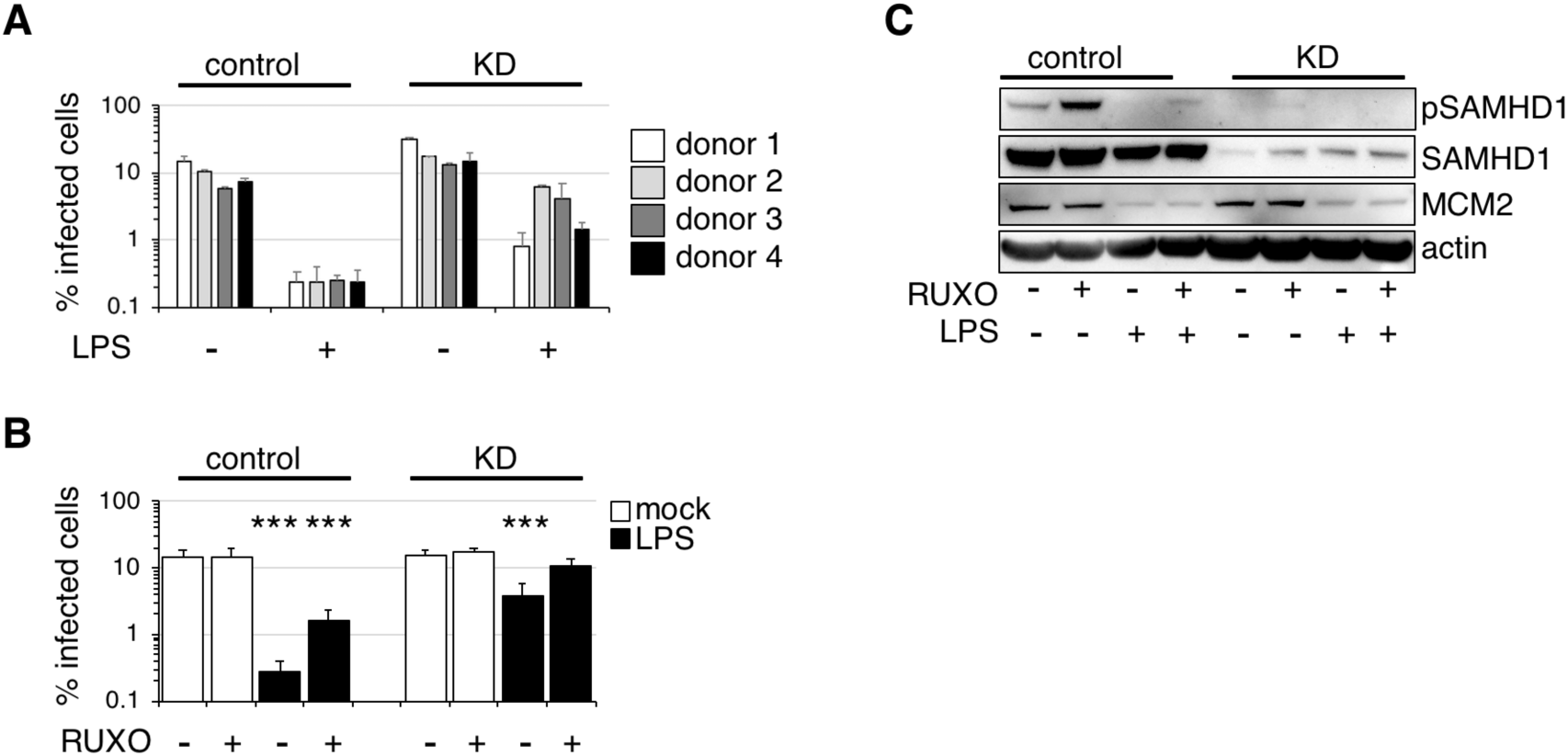
SAMHD1 depletion rescues HIV-1 infection after TLR4 activation. A. MDM were transfected with control or pool of SAMHD1 siRNAs and 3 days later treated with LPS and infected in the presence of LPS with VSV-G-pseudotyped HIV-1 GFP 18h later. The percentage of infected cells was quantified 48h post-infection. Error bars represent technical triplicates. B. MDM were transfected with control or pool of SAMHD1 siRNAs and 3 days later treated with RUXO and 6h later with LPS. Cells were infected in the presence of LPS with VSV-G-pseudotyped HIV-1 GFP 18h later. The percentage of infected cells was quantified 48h post-infection. (*n* = 4, mean ± s.e.m.; ***P-value≤*0.001,* paired *t*-test). C. Cells from a representative donor were used for immunoblotting.

We knocked-down SAMHD1 expression in human MDM using siRNA and infected MDM in the presence or absence of LPS in four different donors (Fig.5A). SAMHD1 KD lifted HIV-1 block in the presence of LPS, though as predicted from data in figure 1, full rescue of infection required addition of RUXO (Fig.5B). Immunoblot confirmed 80% SAMHD1 KD with no effect on the cell cycle marker MCM2 (Fig.5C). We conclude that SAMHD1 plays a key role in the TLR4 mediated antiretroviral state in human macrophages.

## Gram-negative bacteria induce TLR4 activation and G0 arrest in human MDM

The pHrodo™ *E. coli* BioParticles™ are inactivated, unopsonized *E. coli* (K-12 strain) (pHrodo) which function as sensitive, fluorogenic particles for the detection of phagocytic ingestion. We incubated pHrodo with MDM in the presence or absence of different inhibitors for 1h at 37°C (Fig.6A). Unbound pHrodo was washed of and cell incubated overnight when cell supernatants were collected for cytokine detection (Fig.6B) and MDM were infected with HIV-1. Firstly, phagocytosis of pHrodo was unaffected by presence of TLR4, JAK1/2 or TBK1 inhibitors (Fig.6A). Secondly, binding/ingestion of pHrodo triggered expression of TNFa, IL-6 and IL-8 that was abrogated after TLR4 inhibition but not by inhibition of the IFN signalling pathway (Fig.6B). This was confirmed by a IRF3, NFkB translocation assay (Fig.6C). These data clearly show that pHrodo triggers strong immune response in MDM that can be prevented by TLR4 inhibition. Treatment of MDM with pHrodo induced potent HIV-1 inhibition that was accompanied by G0 arrest and SAMHD1 activation/dephosphorylation at T592 (Fig.6D,E)

**FIGURE 6:**
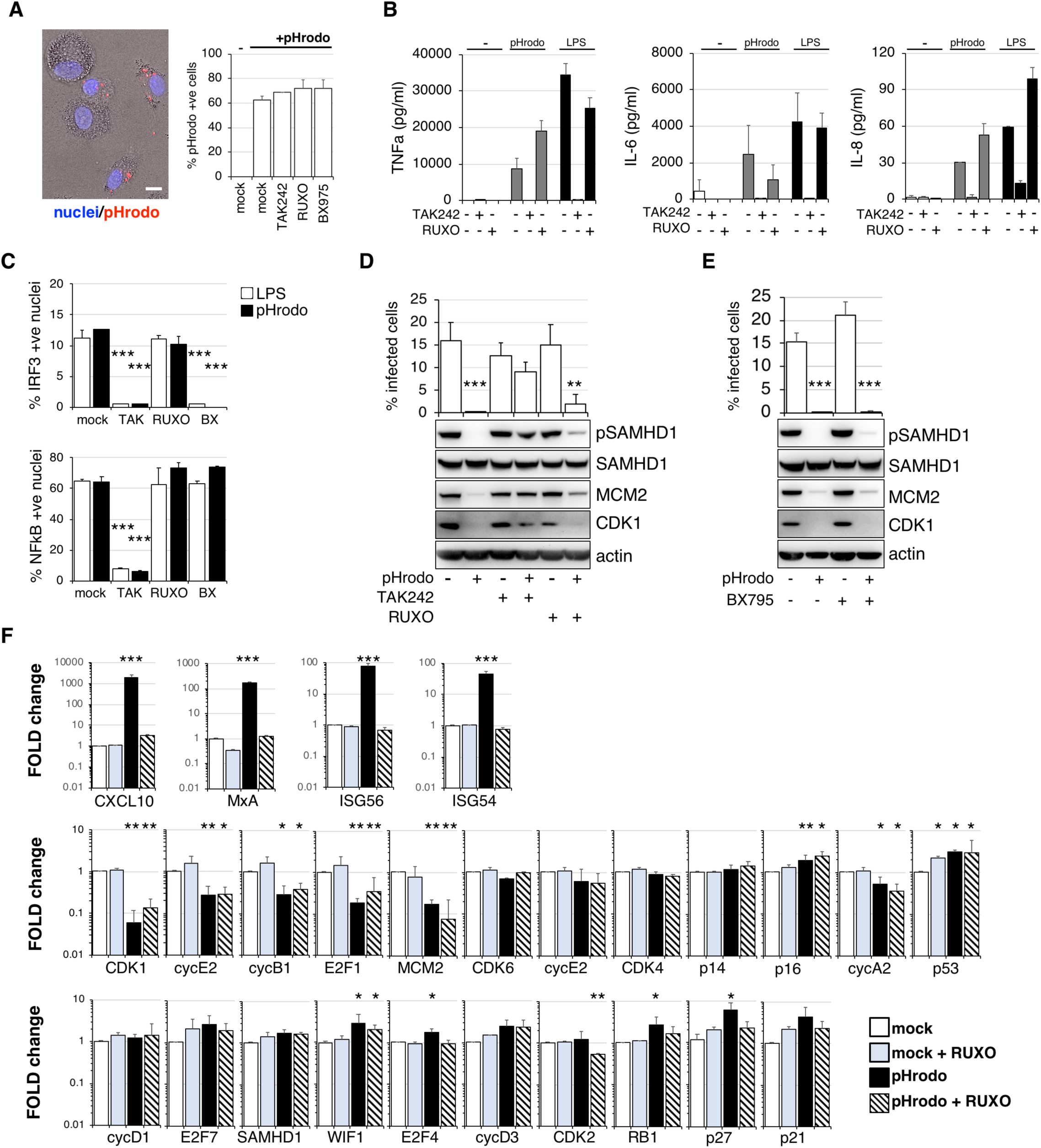
TLR4 activation by gram negative bacteria arrests cell cycle in human MDM and blocks HIV-1 infection. A. pHrodo (a pH-sensitive, rhodamine-based dye)-labeled *E. coli* were added to MDM for 1h. MDM were washed 3x in PBS and fixed. 10^4^ cells were recorded and analysed. Percentage of E.coli positive cells was determined using automated cell-imaging system Hermes WiScan and ImageJ. B. MDM were treated with pHrodo E.coli or LPS in the presence or absence of RUXO and cytokines were measured in culture media 24h later. C. IRF3/NFkB translocation assay. Cells were exposed to pHrodo E.coli in the presence or absence of inhibitors and 2h later stained for IRF3/NFkB. % of cells with nuclear staining was determined. (*n* = 3, mean ± s.e.m.; ***P-value≤*0.00*1, paired *t*-test). MDM were treated with TAK242 or RUXO 6h before addition of pHrodo E.coli. D. Cells were infected by VSV-G pseudotyped HIV-1 18h later. % infected cells were determined 48h post-infection. Cells from a representative donor were used for immunoblotting. (*n* = 3, mean ± s.e.m.; ***P-value≤*0.001*,*\*\*P*-value ≤ 0.01, paired *t*-test). E. MDM were treated with BX795 2h before addition of pHrodo E.coli. Cells were infected by VSV-G pseudotyped HIV-1 18h later. % infected cells were determined 48h post-infection. Cells from a representative donor were used for immunoblotting. (*n* = 3, mean ± s.e.m.; ***P-value≤*0.00*1, paired *t*-test). F. Relative expression levels (fold changes) of cell cycle associated transcripts. MDM were treated with RUXO 6h before addition of pHrodo E.coli. Cells were collected 24h later. (*n* = 3, mean ± s.e.m.; ***P-value≤*0.001, **P*-value ≤ 0.01, *\*P*-value ≤ 0.1, paired *t*-test).

Importantly, TLR4 blockade was able to rescue cell cycle changes/SAMHD1 phosphorylation and HIV-1 infection but neither RUXO or TBK1 inhibitor BX795 could restore infection and cell cycle changes, phenocopying experiments with LPS alone (Fig.2). However, when we measured cell cycle associated transcripts (Fig.6F) there were several significant differences between LPS and E. coli mediated TLR4 activation. Firstly, p16 was significantly increased and cycA2 decreased. Both these transcripts were unchanged after IFN signalling inhibition suggesting they are IFN-independent changes (Fig.6F, compare to Fig.3 and 4). These data are the first evidence of G0 arrest in human macrophages after gram-negative bacterial exposure. This G0 arrest is IFN independent and generates an anti-viral environment.

## Discussion

Here we have reported three key novel observations. Firstly, G0 arrest occurs in primary human macrophages after exposure to pathogen via TLR4. Secondly, it is MyD88 and thus an NFkB independent phenomenon. Thirdly, G0 arrest is mediated through two independent pathways, one of which is IFN-dependent and the other IFN-independent.

Our previous work had shown that human monocyte derived macrophages (MDM) can re-enter cell cycle from their G0 state ^19, 21^. Once the cells enter cell cycle they are permissive to HIV-1 infection. We took advantage of this experimental model to investigate whether the cell cycle is involved in HIV-1 restriction after TLR4 activation. Indeed, TLR4 activation caused G0 arrest that correlated with SAMHD1 activation and HIV-1 restriction in macrophages. Moreover, this arrest and subsequent blockade to HIV-1 infection cannot be simply overcome by blocking IFN signalling. Demonstrating that IFN is not the major driver, but one of two independent mediators of HIV-1 restriction after TLR4 activation is an important finding.

The effect of LPS on G0 arrest has been shown in mouse primary cells and murine cell lines ^31, 32^ or in the human cell line, THP-1/U937 ^33, 34^. Despite these reports, surprisingly little is known about the mechanism how LPS causes G0 arrest. It has been suggested that production of ROS or DNA damage can play role ^34, 35^. Our data suggest that neither ROS nor DNA damage seems to be responsible for G0 arrest in human MDM (Fig.S2, S3).

We speculate that mitogen activated protein kinase kinases may play role in the mechanism. We have shown previously that cell cycle re-entry from G0 to G1 state is mediated by MEK/ERK kinases. These kinases can be activated by TLR4 activation in a MyD88-dependent and independent manner ^36, 37^. Even though it might seem to be counter intuitive as our previous observations show that activation of MEK/ERK leads to cell cycle re-entry not to G0 arrest in MDM ^19^, it has been suggested that prolonged exposure to LPS can downregulate phosphorylation (activation) of these kinases ^38^, possibly leading thus to G0 transition. Another possibility is that strong TLR4 activation triggers an apoptotic program in the cells and G0 arrest is first step to apoptosis and cell death ^39^. However, we do not see extensive reduction in cell numbers even 5 days post-LPS treatment (Fig.S3D), suggesting survival of activated macrophages.

As with the use of LPS, the majority of studies investigating the effect of IFN on G0 arrest have been performed in murine cells and transformed cell lines ^27–29^. Very little is known about IFN regulation of cell cycle in non-malignant cells. We show here that human primary macrophages exit the cell cycle after exposure to exogenous IFNβ) and that this effect is dependent on JAK/STAT signalling. It has been shown that SAMHD1 can be activated/dephosphorylated by type I,II and III IFN ^16, 22^ in macrophages. We confirmed this observation and showed that SAMHD1 phosphorylation and HIV-1 infection can be rescued when JAK/STAT signalling is blocked after addition of exogenous IFNβ). Nevertheless, after activation of the TLR4 receptor, SAMHD1 remains active/dephosphorylated even in the presence of IFN/JAK/STAT signalling inhibitor, highlighting the role of two independent pathways regulating HIV-1 restriction in LPS activated macrophages.

The interferon-independent pathway is accompanied by p21 upregulation and SAMHD1 dephosphorylation. SAMHD1 knockdown (KD) substantially relieved the block to HIV-1 infection after LPS addition. However, only a combination of SAMHD1 KD and inhibition of IFN signalling could achieve complete rescue of HIV-1 infection after TLR4 activation. This is not surprising as many IFN-inducible proteins with HIV-1 restriction potency have been shown ^40^ and may play additional roles in this restriction. These data confirm that SAMHD1 is major player in LPS mediated HIV-1 restriction.

Importantly we have also shown that E. coli leads to G0 arrest in human macrophages and this arrest cannot be rescued by blocking IFN signalling. Remarkably however, inhibition of TLR4 was able to prevent the G0 arrest and SAMHD1 phosphorylation. This suggests that whole gram-negative bacteria also activate two independent pathways to induce G0 arrest.

Why would macrophages regulate their cell cycle? Macrophages contribute to innate immunity via phagocytosis, antigen presentation but they are also secretory cells vital to the regulation of immune responses and development of inflammation. One can imagine that cell division of macrophages would benefit the host by increasing the number of effector cells at the centre of infection. But at the same time the division of infected cells harbouring live pathogen could also lead to doubling of infected cells, an event that can potentially harm the host. Even though our previous work showed that MDM re-enter cell cycle without measurable cell division ^19^, many tissue resident macrophages can proliferate ^41^ and thus could use G0 arrest as a response to danger signal to stop division and contain infection.

It is also possible that cell cycle changes are necessary for activation of non-cycling function of cell cycle associated proteins. G0 or G0 arrested cells will increase expression of e.g. p14, p16, p21 or p27 proteins. It has been shown that cell cycle regulators can serve non-cycling functions in innate immunity. It has been suggested that CDK activity is required for IFN-b production ^42^, which in turn initiates immune system activation. Nevertheless, these experiments still await validation by specific CDK knockdowns. It has been shown that p21 supress IL-1b ^43^, or that p16 inhibits macrophage activity by degradation of interlukin-1 receptor and thus impairs IL-6 production that can lead to tissue inflammation reduction ^44^. Moreover, p27 seems to have a unique role in macrophage migration ^45^. It is thus conceivable that cell cycle regulators can contribute to maintenance of balanced responses to immune stimuli.

The concept that immune system can be manipulated by the host cell cycle has therapeutic implications and establishes a new paradigm for understanding not only basic cell biology but may present new ways to treat infectious disease but possibly also autoimmune diseases and cancer.

The relevance of macrophage G0 arrest by LPS may be in relevant in HIV pathogenesis where macrophages may be exposed to gut derived LPS during inflammation in the acute or chronic phase of HIV. These macrophages are likely arrested with SAMHD1 dephosphorylated at T592, and thereby rendered resistant to HIV-1. T cells would remain more susceptible as they do not express TLR4 and are less sensitive to IFN, consistent with observed high levels of viral turnover in GALT associated CD4 T cells.

In summary, our data show that TLR4 activation regulates the cell cycle in human primary macrophages after exposure to immune stimuli or pathogen. We also show evidence that TLR4 activation by bacterial LPS can mediate HIV-1 inhibition through regulation of SAMHD1 phosphorylation in a MyD88-independent manner by two pathways: (i) IFN dependent pathway and (ii) IFN-independent pathway accompanied by p21 upregulation and SAMHD1 dephosphorylation. Together, these data suggest that macrophages can rapidly achieve an anti-viral state by activation of TLR4 and G0 arrest prior to IFN secretion, thereby demonstrating redundancy and highlighting the importance of cell cycle regulation as a response to danger signals. Finally, interferon independent activation of SAMHD1 by TLR4 represents a novel mechanism for limiting the HIV-1 reservoir size and should be considered for host-directed therapeutic approaches that may contribute to curative interventions.

## Methods

### Reagents, inhibitors, antibodies, plasmids

Tissue culture media and supplements were obtained from Invitrogen (Paisley, UK), and tissue culture plastic was purchased from TPP (Trasadingen, Switzerland). FCS (FBS) was purchased from Biosera (Boussens, France) and Sigma (Sigma, St. Louis, USA). Human serum from human male AB plasma was of USA origin and sterile-filtered (Sigma). All chemicals, were purchased from Sigma (St. Louis, MO, USA) unless indicated otherwise. LPS (Insight Biotechnology, UK), Interferon-beta (PeproTech, UK), CellRox (Invitrogen, UK), Tenascin-C (Bio-Techne, Minneapolis, MN, USA), E.coli pHrodo Bioparticles (ThermoFisher scientific, UK). Ruxolitinib (Cambridge Bioscience, UK), BX795 (Generon, UK), TAK242 (Millipore, UK), 1400W (2B Scientific, UK). Antibodies used were: anti-cdc2 (Cell Signaling Technology, Beverly, USA); anti-SAMHD1 (ab67820, Abcam, UK), beta-actin (ab6276, abcam, UK); mouse anti-MCM2 (BM-28, BD Biosciences, UK); pSAMHD1 ProSci (Poway, CA, USA); p21(sc-6246, Santa Cruz Biotechnology); IRF3 (11904P, Cell Signaling Technology); NFkB p65 (F-6, Insight Biotechnology, UK), gH2AX (613402, BioLegend); 53BP1 (612522, BD Biosciences, UK), Anti-IFNα/β Receptor (PBL Interferon Source), IgG2A antibody (R&D systems, Minneapolis, MN, USA). Anti-TNFa, anti-IL6, anti-IL8, anti-CXCL10 were purchased from BD Biosciences, UK.

### Cell lines and viruses

293T cells were cultured in DMEM complete (DMEM supplemented with 100 U/ml penicillin, 0.1 mg/ml streptomycin, and 10% FCS). VSV-G HIV-1 GFP virus was produced by transfection of 293T with GFP-encoding genome CSGW, packaging plasmid p8.91 and pMDG as previously described ^46^.

### Monocyte isolation and differentiation

PBMC were prepared from HIV seronegative donors (after informed consent was obtained), by density-gradient centrifugation (Lymphoprep, Axis-Shield, UK). Monocyte-derived macrophages (MDM) were prepared by adherence with washing of non-adherent cells after 2h, with subsequent maintenance of adherent cells in RPMI 1640 medium supplemented with 10% human serum and MCSF (10ng/ml) for 3 days and then differentiated for a further 4 days in RPMI 1640 medium supplemented with 10% fetal calf sera without M-CSF.

### Infection of primary cells using full-length and VSV-G pseudotyped HIV-1 viruses

GFP containing VSV-G pseudotyped HIV-1 was added to MDM and after 4h incubation removed and cells were washed in culture medium. The percentage of infected cells was determined 48h post-infection by Hermes WiScan automated cell-imaging system (IDEA Bio-Medical Ltd. Rehovot, Israel) and analysed using MetaMorph and ImageJ software.

### SDS-PAGE and Immunoblots

Cells were lysed in reducing Laemmli SDS sample buffer containing PhosSTOP (Phosphatase Inhibitor Cocktail Tablets, Roche, Switzerland) at 96°C for 10 minutes and the proteins separated on NuPAGE® Novex® 4-12% Bis-Tris Gels. Subsequently, the proteins were transferred onto PVDF membranes (Millipore, Billerica, MA, USA), the membranes were quenched, and proteins detected using specific antibodies. Labelled protein bands were detected using Amersham ECL Prime Western Blotting Detection Reagent (GE Healthcare, USA) and Amersham Hyperfilm or AlphaInnotech CCD camera. Protein band intensities were recorded and quantified using AlphaInnotech CCD camera and AlphaView software (ProteinSimple, San Jose, California, USA).

### SAMHD1 knock-down by siRNA

1x10e5 MDM differentiated in MCSF for 4 days were transfected with 20pmol of siRNA (L-013950-01, Dharmacon) using Lipofectamine RNAiMAX Transfection Reagent (Invitrogen). Transfection medium was replaced after 18h with RPMI 1640 medium supplemented with 10% FCS and cells cultured for additional 3 days before infection.

### Quantitative PCR

Total RNA was isolated from macrophages using the Total RNA Purification Kit from Norgen Biotek (Thorold, Canada). cDNA was synthesised using Superscript III Reverse Transcriptase (Thermo Fisher Scientific) using 500ng of template RNA. qPCR was performed on ABI 7300 machine (Thermo Fisher Scientific) using Fast SYRB green master mix (Thermo Fisher Scientific). Expression levels of target genes were normalised to glyceraldehyde-3-phosphate dehydrogenase (GAPDH) as previously described ^47^. See primer sequences in supplementary Table1.

### Immunofluorescence

MDMs were fixed in 3% PFA, quenched with 50 mM NH_4_Cl and permeabilized with 0.1% Triton X-100 in PBS or 90% Methanol. After blocking in PBS/1% FCS, MDMs were labelled for 1 hour with primary antibodies diluted in PBS/1% FCS, washed and labelled again with Alexa Fluor secondary antibodies for 1 hour. Cells were washed in PBS/1% FCS and stained with DAPI in PBS for 20 minutes. Labelled cells were detected using Hermes WiScan automated cell-imaging system (IDEA Bio-Medical Ltd. Rehovot, Israel) and analysed using MetaMorph and ImageJ software.

### Phagocytosis assay using pHrodo Bioparticles

MDM were exposed to 0.25ug pHrodo (a pH-sensitive, rhodamine-based dye)-labeled *E. coli* for 1h. MDM were washed 3x in PBS and fixed. Percentage of E.coli positive cells was determined using Hermes WiScan automated cell-imaging system (IDEA Bio-Medical Ltd. Rehovot, Israel) and analysed using MetaMorph and ImageJ software. 10^4^ cells were recorded and analysed.

## ELISA

Medium was collected and cytokines levels detected by ELISA (BD Biosciences) according to the manufacturer’s instructions.

## Acknowledgments

This work was funded by a Wellcome Trust Senior Fellowship in Clinical Science to RKG (WT108082AIA) and the National Institute for Health Research University College London Hospitals Biomedical Research Centre.

## Ethics Statement

Adult subjects provided written informed consent. Primary Macrophage & Dendritic Cell Cultures from Healthy Volunteer Blood Donors has been reviewed and granted ethical permission by the National Research Ethics Service through The Joint UCL/UCLH Committees on the Ethics of Human Research (Committee Alpha) 2nd of December 2009. Reference number 06/Q0502/92.

## Author contributions

PM, RKG, LZA designed experiments; PM, RKG wrote the manuscript; PM, HW performed experiments; PM, RKG, LZA analysed data.

## SUPPLEMENTARY FIGURES

**Figure S1.**
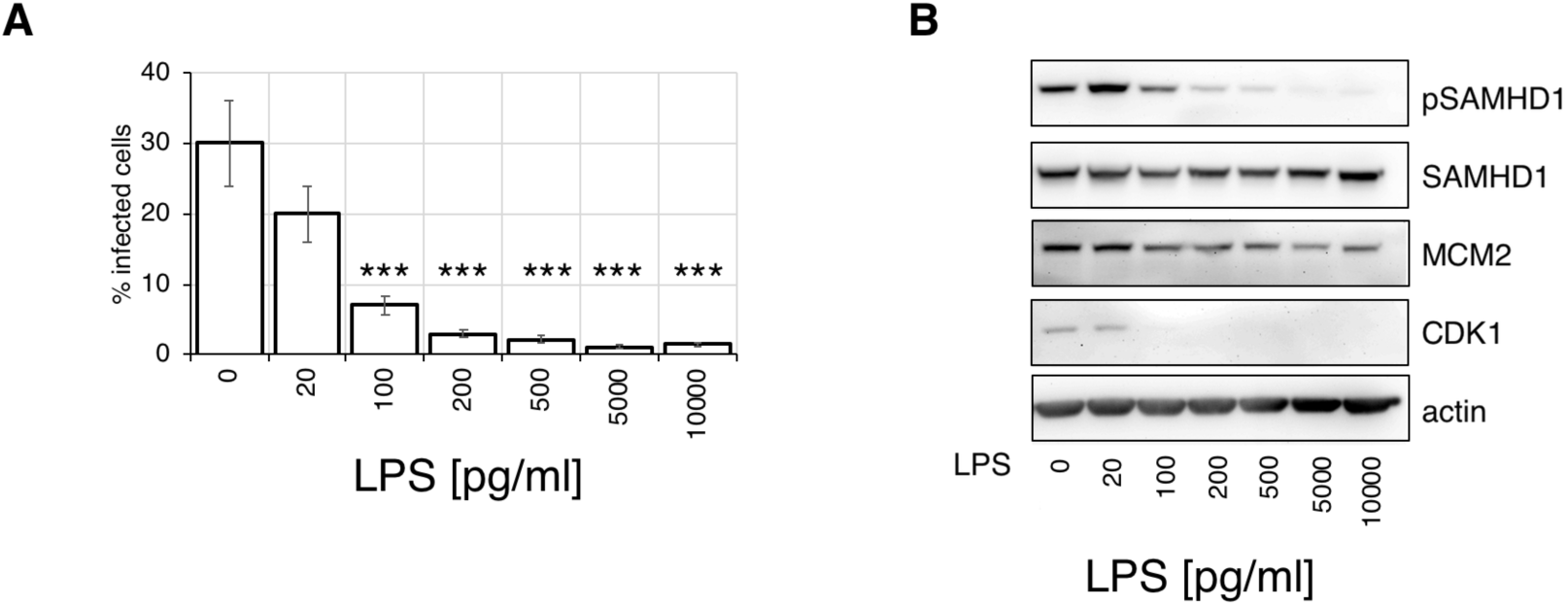
**A.** MDM were treated with increasing concentration of LPS 18h before infection. Cells were infected by VSV-G pseudotyped HIV-1 and % infected cells were determined 48h post-infection. (*n* = 3, mean ± s.e.m.; ***P-value≤*0.001,* paired *t*-test). **B.** Cells from a representative donor were used for immunoblotting.

**Figure S2.**
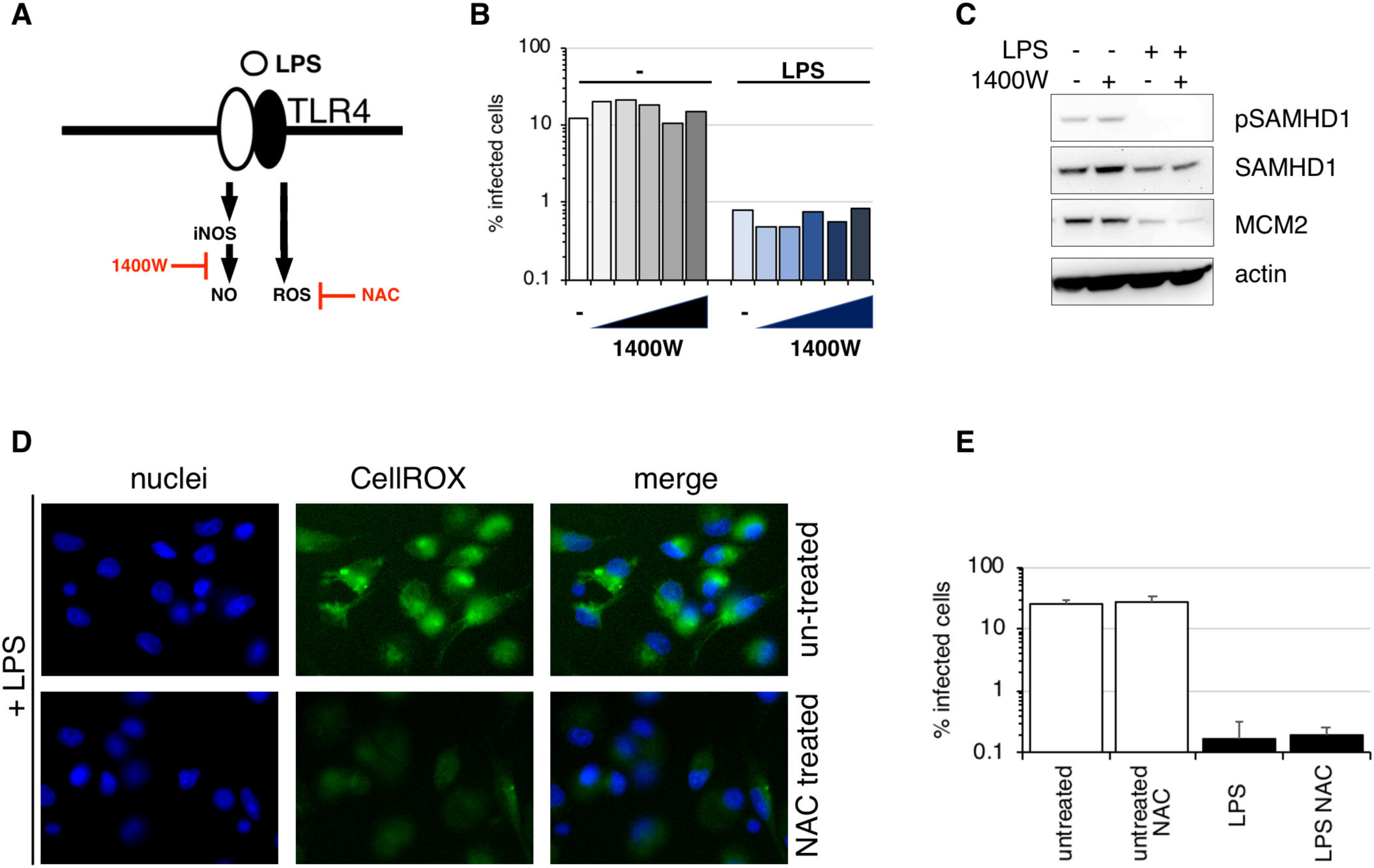
A. Simplified diagram of site of function of iNOS and ROS inhibitors. B. MDM were treated with increasing concentration of iNOS inhibitor 1400W 6h before LPS addition. LPS was added to cells 18h before infection. Cells were infected by VSV-G pseudotyped HIV-1 and % infected cells were determined 48h post-infection. C. Cells from a representative donor were used for immunoblotting. D. MDM were treated with LPS in the presence or absence of ROS inhibitor NAC and labelled with CellROX to detect ROS. E. MDM were treated with LPS in the presence or absence of ROS inhibitor NAC 18h before infection. Cells were infected by VSV-G pseudotyped HIV-1 and % infected cells were determined 48h post-infection. (*n* = 3, mean ± s.e.m.).

**Figure S3.**
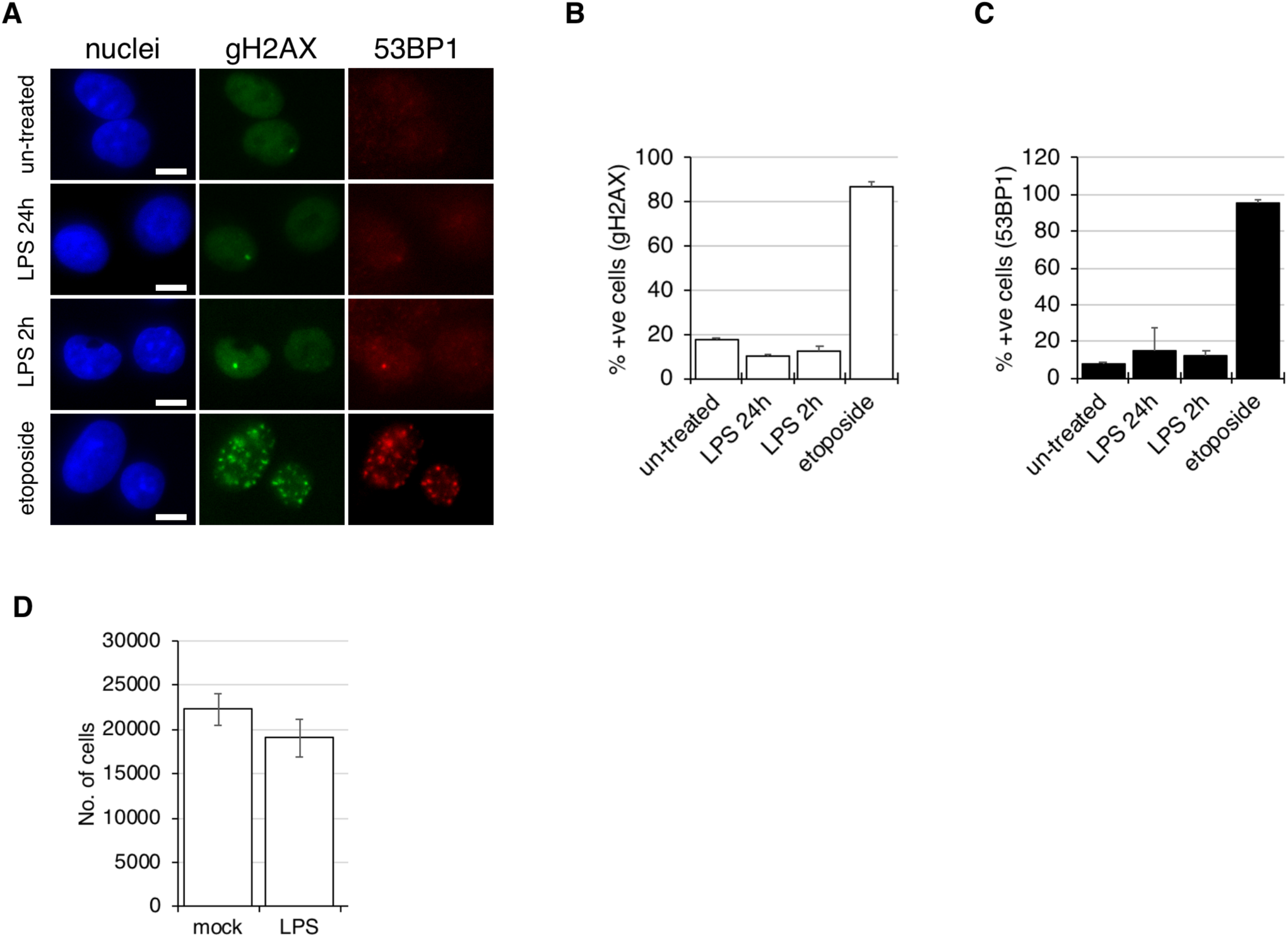
A. MDM were treated with LPS for 2 or 24h, or with Etoposide for 2h (positive control for DNA damage). Cells were fixed and stained for DNA damage foci positive for gH2AX and 53BP1. B. Quantification of gH2AX positive cells. 1,000 cells were analysed using Hermes WiScan automated cell-imaging system (IDEA Bio-Medical Ltd. Rehovot, Israel) and analysed using MetaMorph and ImageJ software. C. Quantification of 53BP1 positive cells. 1,000 cells were analysed using Hermes WiScan automated cell-imaging system (IDEA Bio-Medical Ltd. Rehovot, Israel) and analysed using MetaMorph and ImageJ software. D. MDM treated or untreated with LPS were stained for nuclei using DAPI stain. Cell numbers were quantified using Hermes WiScan automated cell-imaging system (IDEA Bio-Medical Ltd. Rehovot, Israel) and analysed using MetaMorph and ImageJ software. (*n* = 7, mean ± s.e.m.; (ns) non-significant, paired *t*-test).

## SUPPLEMENTARY TABLE 1

**Supplementary Table 1.**
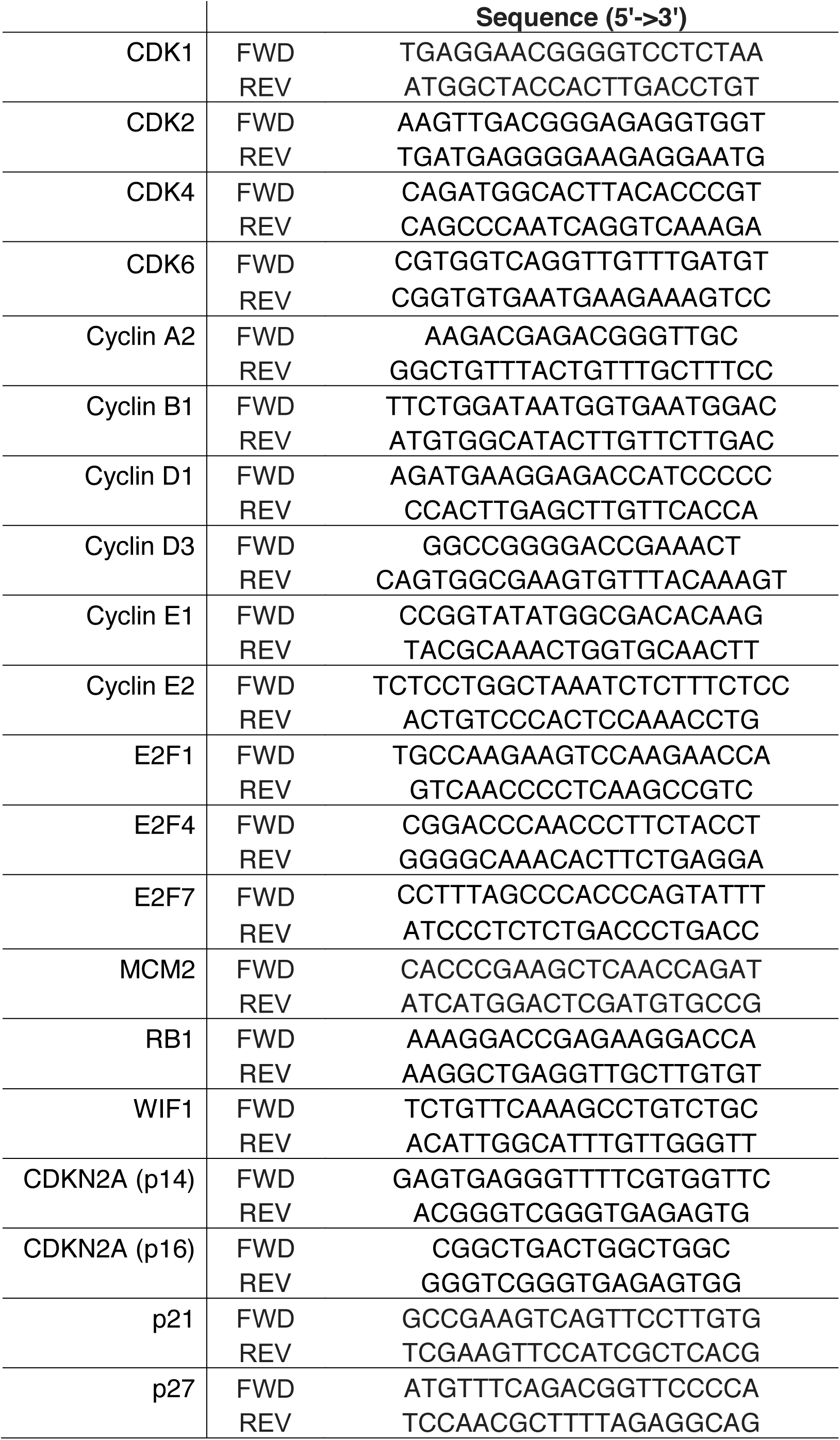
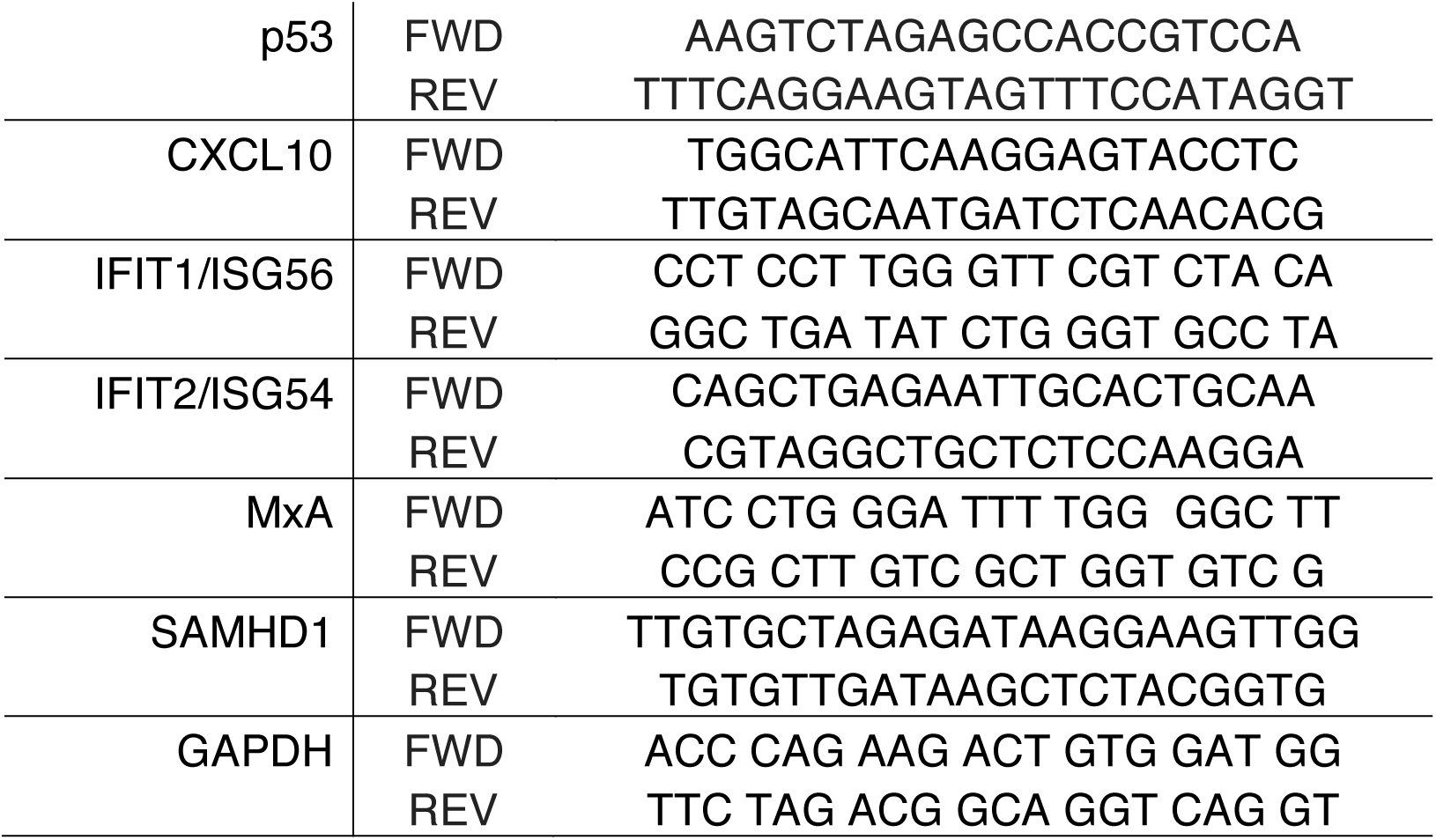
Sequences of primers used for this study.

## References

1. Iwasaki, A. & Medzhitov, R. Toll-like receptor control of the adaptive immune responses. Nature Immunology (2004). doi:10.1038/ni1112

2. Park, B. S. & Lee, J. O. Recognition of lipopolysaccharide pattern by TLR4 complexes. Experimental and Molecular Medicine (2013). doi:10.1038/emm.2013.97

3. Akira, S. & Takeda, K. Toll-like receptor signalling. Nature Reviews Immunology (2004). doi:10.1038/nri1391

4. Geonnotti, A. R. et al. Differential Inhibition of Human Immunodeficiency Virus Type 1 in Peripheral Blood Mononuclear Cells and TZM-bl Cells by Endotoxin-Mediated Chemokine and Gamma Interferon Production. AIDS Res. Hum. Retroviruses (2010). doi:10.1089/aid.2009.0186

5. Franchin, G. et al. Lipopolysaccharide Inhibits HIV-1 Infection of Monocyte-Derived Macrophages Through Direct and Sustained Down-Regulation of CC Chemokine Receptor 5. J. Immunol. (2014). doi:10.4049/jimmunol.164.5.2592

6. Kornbluth, R. S. Interferons and bacterial lipopolysaccharide protect macrophages from productive infection by human immunodeficiency virus in vitro. J. Exp. Med. (2004). doi:10.1084/jem.169.3.1137

7. Schlaepfer, E., Rochat, M.-A., Duo, L. & Speck, R. F. Triggering TLR2, -3, -4, -5, and -8 Reinforces the Restrictive Nature of M1- and M2-Polarized Macrophages to HIV. J. Virol. (2014). doi:10.1128/jvi.01053-14

8. Bernstein, M. S., Tong-Starksen, S. E. & Locksley, R. M. Activation of human monocyte-derived macrophages with lipopolysaccharide decreases human immunodeficiency virus replication in vitro at the level of gene expression. J. Clin. Invest. (1991). doi:10.1172/JCI115337

9. Verani, A., Sironi, F., Siccardi, A. G., Lusso, P. & Vercelli, D. Inhibition of CXCR4-Tropic HIV-1 Infection by Lipopolysaccharide: Evidence of Different Mechanisms in Macrophages and T Lymphocytes. J. Immunol. (2014). doi:10.4049/jimmunol.168.12.6388

10. Reinhard, C., Bottinelli, D., Kim, B. & Luban, J. Vpx rescue of HIV-1 from the antiviral state in mature dendritic cells is independent of the intracellular deoxynucleotide concentration. Retrovirology (2014). doi:10.1186/1742-4690-11-12

11. Verani, B. A. et al. C – C Chemokines Released by Lipopolysaccharide Infection in Both Macrophages and T Cells. 185, (1997).

12. Wang, H., Sun, J. & Goldstein, H. Human Immunodeficiency Virus Type 1 Infection Increases the In Vivo Capacity of Peripheral Monocytes To Cross the Blood-Brain Barrier into the Brain and the In Vivo Sensitivity of the Blood-Brain Barrier to Disruption by Lipopolysaccharide. J. Virol. (2008). doi:10.1128/JVI.00768-08

13. Mehta, H. V., Jones, P. H., Weiss, J. P. & Okeoma, C. M. IFN-and Lipopolysaccharide Upregulate APOBEC3 mRNA through Different Signaling Pathways. J. Immunol. (2012). doi:10.4049/jimmunol.1200777

14. Goldstone, D. C. et al. HIV-1 restriction factor SAMHD1 is a deoxynucleoside triphosphate triphosphohydrolase. Nature (2011). doi:10.1038/nature10623

15. Lahouassa, H. et al. SAMHD1 restricts the replication of human immunodeficiency virus type 1 by depleting the intracellular pool of deoxynucleoside triphosphates. Nat. Immunol. (2012). doi:10.1038/ni.2236

16. Cribier, A., Descours, B., Valadão, A. L. C., Laguette, N. & Benkirane, M. Phosphorylation of SAMHD1 by Cyclin A2/CDK1 Regulates Its Restriction Activity toward HIV-1. Cell Rep. (2013). doi:10.1016/j.celrep.2013.03.017

17. White, T. E. et al. The Retroviral Restriction Ability of SAMHD1, but Not Its Deoxynucleotide Triphosphohydrolase Activity, Is Regulated by Phosphorylation. Cell Host Microbe 13, 441–451 (2013).

18. Arnold, L. H. et al. Phospho-dependent Regulation of SAMHD1 Oligomerisation Couples Catalysis and Restriction. PLoS Pathog. (2015). doi:10.1371/journal.ppat.1005194

19. Mlcochova, P. et al. A G1-like state allows HIV-1 to bypass SAMHD1 restriction in macrophages. EMBO J. (2017). doi:10.15252/embj.201696025

20. Mlcochova, P., Caswell, S. J., Taylor, I. A., Towers, G. J. & Gupta, R. K. DNA damage induced by topoisomerase inhibitors activates SAMHD1 and blocks HIV-1 infection of macrophages. EMBO J. (2017). doi:10.15252/embj.201796880

21. Mlcochova, P., Caswell, S. J., Taylor, I. A., Towers, G. J. & Gupta, R. K. DNA damage induced by topoisomerase inhibitors activates SAMHD1 and blocks HIV-1 infection of macrophages. EMBO J. 37, (2018).

22. Szaniawski, M. A. et al. SAMHD1 Phosphorylation Coordinates the Anti-HIV-1 Response by Diverse Interferons and Tyrosine Kinase Inhibition. MBio (2018). doi:10.1128/mbio.00819-18

23. Ajayi, S. et al. Ruxolitinib. in *Recent Results in Cancer Research* (2018). doi:10.1007/978-3-319-91439-8_6

24. Clark, K., Plater, L., Peggie, M. & Cohen, P. Use of the pharmacological inhibitor BX795 to study the regulation and physiological roles of TBK1 and IκB Kinase ∈:A distinct upstream kinase mediates ser-172 phosphorylation and activation. J. Biol. Chem. (2009). doi:10.1074/jbc.M109.000414

25. Midwood, K. S., Chiquet, M., Tucker, R. P. & Orend, G. Tenascin-C at a glance. J. Cell Sci. (2016). doi:10.1242/jcs.190546

26. Midwood, K. et al. Tenascin-C is an endogenous activator of Toll-like receptor 4 that is essential for maintaining inflammation in arthritic joint disease. Nat. Med. (2009). doi:10.1038/nm.1987

27. Xaus, J. et al. Interferon γ induces the expression of p21(waf-1) and arrests macrophage cell cycle, preventing induction of apoptosis. Immunity (1999). doi:10.1016/S1074-7613(00)80085-0

28. Dey, A. et al. Colony-stimulating Factor-1 Receptor Utilizes Multiple Signaling Pathways to Induce Cyclin D2 Expression. Mol. Biol. Cell (2013). doi:10.1091/mbc.11.11.3835

29. Zhang, C., Cui, G., Chen, Y. & Fan, K. Antitumor effect of interferon-on U937 human acute leukemia cells in vitro and its molecular mechanism. J. Huazhong Univ. Sci. Technol. -Med. Sci. (2007). doi:10.1007/s11596-007-0509-z

30. Munn, D. H., Pressey, J., Beall, A. C., Hudes, R. & Alderson, M. R. Selective activation-induced apoptosis of peripheral T cells imposed by macrophages. A potential mechanism of antigen-specific peripheral lymphocyte deletion. J Immunol (1996).

31. Vairo, G., Royston, A. K. & Hamilton, J. A. Biochemical events accompanying macrophage activation and the inhibition of colony-stimulating factor-1-induced macrophage proliferation by tumor necrosis factor-α, interferon-γ, and lipopolysaccharide. J. Cell. Physiol. (1992). doi:10.1002/jcp.1041510324

32. Zhang, K. et al. MicroRNA-322 inhibits inflammatory cytokine expression and promotes cell proliferation in LPS-stimulated murine macrophages by targeting NF- κB1 (p50). Biosci. Rep. (2016). doi:10.1042/bsr20160239

33. Thongngarm, T., Jenkins, J. K., Ndebele, K. & McMurray, R. W. Estrogen and progesterone modulate monocyte cell cycle progression and apoptosis. Am. J. Reprod. Immunol. (2003). doi:10.1034/j.1600-0897.2003.00015.x

34. Mytych, J., Romerowicz-Misielak, M. & Koziorowski, M. Long-term culture with lipopolysaccharide induces dose-dependent cytostatic and cytotoxic effects in THP-1 monocytes. Toxicol. Vitr. (2017). doi:10.1016/j.tiv.2017.03.009

35. Sagar, S., Kumar, P., Behera, R. R. & Pal, A. Effects of CEES and LPS synergistically stimulate oxidative stress inactivates OGG1 signaling in macrophage cells. J. Hazard. Mater. (2014). doi:10.1016/j.jhazmat.2014.05.096

36. Lee, S. H. et al. ERK activation drives intestinal tumorigenesis in Apc min/+ mice. Nat. Med. (2010). doi:10.1038/nm.2143

37. Kawai, T. et al. Lipopolysaccharide stimulates the MyD88-independent pathway and results in activation of IFN-regulatory factor 3 and the expression of a subset of lipopolysaccharide-inducible genes. J. Immunol. (2001).

38. Perkins, D. J., Qureshi, N. & Vogel, S. N. A Toll-Like Receptor-Responsive Kinase, Protein Kinase R, Is Inactivated in Endotoxin Tolerance through Differential K63/K48 Ubiquitination. MBio (2010). doi:10.1128/mbio.00239-10

39. Evan, G. I. & Vousden, K. H. Proliferation, cell cycle and apoptosis in cancer. Nature (2001). doi:10.1038/35077213

40. Neil, S. & Bieniasz, P. Human immunodeficiency virus, restriction factors, and interferon. J. Interferon Cytokine Res. 29, 569–80 (2009).

41. Gomez Perdiguero, E., Schulz, C. & Geissmann, F. Development and homeostasis of ‘resident’ myeloid cells: The case of the microglia. Glia (2013). doi:10.1002/glia.22393

42. Cingöz, O. & Goff, S. P. Cyclin-dependent kinase activity is required for type I interferon production. Proc. Natl. Acad. Sci. (2018). doi:10.1073/pnas.1720431115

43. Scatizzi, J. C. et al. The CDK domain of p21 is a suppressor of IL-1β-mediated inflammation in activated macrophages. Eur. J. Immunol. (2009). doi:10.1002/eji.200838683

44. Murakami, Y., Mizoguchi, F., Saito, T., Miyasaka, N. & Kohsaka, H. p16INK4a Exerts an Anti-Inflammatory Effect through Accelerated IRAK1 Degradation in Macrophages. J. Immunol. (2012). doi:10.4049/jimmunol.1103156

45. Gui, P. et al. Rho/ROCK pathway inhibition by the CDK inhibitor p27kip1 participates in the onset of macrophage 3D-mesenchymal migration. J. Cell Sci. (2014). doi:10.1242/jcs.150987

46. Besnier, C., Takeuchi, Y. & Towers, G. Restriction of lentivirus in monkeys. Proc. Natl. Acad. Sci. (2002). doi:10.1073/pnas.172384599

47. Tsang, J. et al. HIV-1 infection of macrophages is dependent on evasion of innate immune cellular activation. AIDS (2009). doi:10.1097/QAD.0b013e328331a4ce

